# Tissue-specific transcriptomic responses to early *Toxoplasma gondii* infection in sheep during congenital toxoplasmosis

**DOI:** 10.64898/2026.10.27.762155

**Authors:** Montserrat Coronado, Roberto Sánchez-Sánchez, Armando Reyes-Palomares, Natalia Velasco-Jiménez, Marta Silva, Julio Benavides, Luis Miguel Ortega-Mora, Pilar Horcajo

**Author notes:** Correspondence (LMOM) SALUVET Group Animal Health Department Faculty of Veterinary Sciences Complutense University of Madrid Ciudad Universitaria s/n Madrid 28040 (Spain). These authors have contributed equally to this work (Luis Miguel Ortega-Mora, Pilar Horcajo).

## Abstract

*Toxoplasma gondii* is a widespread zoonotic pathogen of major One Health concern, capable of causing congenital toxoplasmosis in both humans and sheep. Despite its impact, early host–parasite molecular interactions during acute infection in pregnant hosts remain largely unexplored. In this study, we characterized the transcriptomic responses of key tissues involved in early infection (the small intestine, mesenteric lymph nodes, and placenta) in pregnant ewes experimentally infected with *T. gondii* strain TgShSp1. RNA-seq analysis revealed pronounced, tissue-specific transcriptional changes at 6 days post-infection (dpi), particularly in the mesenteric lymph nodes, characterized by activation of interferon- and cytokine-mediated immune pathways. In contrast, transcriptional responses at 3 dpi were limited, consistent with a silent phase of parasite dissemination. Despite the absence of detectable parasite DNA, placentomes showed modest interferon-associated responses and upregulation of DNA repair and cellular stress pathways. These changes coincided with marked suppression of cell-cycle progression and extracellular matrix remodeling alongside inhibition of angiogenesis and hypoxia pathways. Altogether, early disruption of placental vascular function and tissue remodeling may contribute to fetal loss. In summary, these findings provide new insights into early host responses to *T. gondii* infection in a pregnant host and identify molecular mechanisms that may predispose to early fetal death during ovine congenital toxoplasmosis.

**IMPORTANCE:** Within a One Health framework, *T. gondii* represents a critical link between human and animal health, as congenital infection can cause reproductive failure, miscarriage, and fetal abnormalities in both humans and livestock. Although the pathogenesis of classical ovine toxoplasmosis is well characterized, abortion occurring during the acute phase of infection without detectable parasite presence in placental or fetal tissues remains poorly understood. This suggests that pregnancy loss may be driven by host-mediated immune and vascular disruptions. Understanding these processes is essential to clarify the pathogenesis of early reproductive failure. By applying transcriptomic approaches to target tissues involved in infection and pregnancy, this study highlights novel molecular pathways and host responses potentially associated with early abortion. Furthermore, as current diagnostic techniques cannot rule out the occurrence of such early events in human congenital toxoplasmosis, this ovine model provides an essential translational tool to explore placental injury and reproductive failure in humans.

## INTRODUCTION

*Toxoplasma gondii* is an obligate intracellular parasite capable of infecting a wide range of warm-blooded animals, including humans and livestock representing a major parasite of global One Health concern. In sheep, *T. gondii* is a major cause of reproductive failure, leading to abortions, stillbirths, and the birth of weak offspring, collectively imposing considerable economic burdens on the global sheep industry (1–3).

Classically, abortions in ovine toxoplasmosis occur from 28 days post-infection (dpi) onwards and are associated with necrotic lesions resulting from extensive parasite multiplication in the placenta and fetal tissues (4). However, experimental studies have reported earlier abortions, occurring between 7 and 14 dpi, characterized by thrombotic lesions and infarcts in placentomes despite the absence of detectable parasites in placental and fetal tissues (5–8). An excessive inflammatory response triggered during early infection has been proposed as a potential driver of vascular injury and thrombosis (4). However, the molecular interactions between the host and parasite in the tissues involved in early infection and pregnancy maintenance remain poorly understood.

Following oral ingestion of sporulated oocysts, *T. gondii* invades the small intestinal epithelium and differentiates into rapidly replicating tachyzoites. The parasite then disseminates through lymphatic and blood circulation. Mesenteric lymph nodes (MLNs) are important sites of early immune activation and may contribute to regulating parasite dissemination (9, 10). In pregnant sheep, the placenta is a target organ where the parasite becomes detectable from 14 dpi (11, 12). Most studies of early infection dynamics and host immune responses have used murine models (10, 13–16). Nevertheless, significant anatomical and physiological differences, especially in the gastrointestinal system (17–19), as well as species-specific variations in the immune response to *T. gondii* infection (20), limit the direct extrapolation of murine data to ovine hosts.

In a recent study, we characterized early *T. gondii* infection in pregnant sheep at 3 and 6 dpi (8). The jejunum and jejunal MLNs emerged as key sites of early parasite replication and dissemination. Although parasite DNA was not detected in placentomes (PL), endothelial activation and thrombotic lesions were observed at 6 dpi, preceding early fetal death at 8 dpi. These findings suggest that placental pathology may develop before detectable parasite colonization.

Here, we aimed to investigate the transcriptomic landscape in these three strategic tissues of early *T. gondii* infection: the intestine, MLN, and PL of the pregnant ewes experimentally infected with *T. gondii*. By analyzing the differential gene expression profiles within these key tissues, our primary objective is to gain a deeper understanding of the molecular mechanisms underlying the pathogenesis of ovine toxoplasmosis, with a particular focus on mechanisms that could predispose to early fetal death. Importantly, within a One Health framework, this ovine model provides a valuable translational tool to address critical gaps in human congenital toxoplasmosis.

## RESULTS

### Transcriptional landscape reveals a conserved proteostasis and stress response signature at 6 dpi following *Toxoplasma gondii*-infection

To elucidate the host response associated with early *T. gondii* infection during gestation, bulk RNA-seq was performed on intestine wall (IW), MLN, and PL tissues obtained from pregnant sheep infected with TgShSp1 and non-infected sheep at early stages of infection (3 and 6 dpi), before the occurrence of fetal death (Fig. 1A).

**Figure 1.**
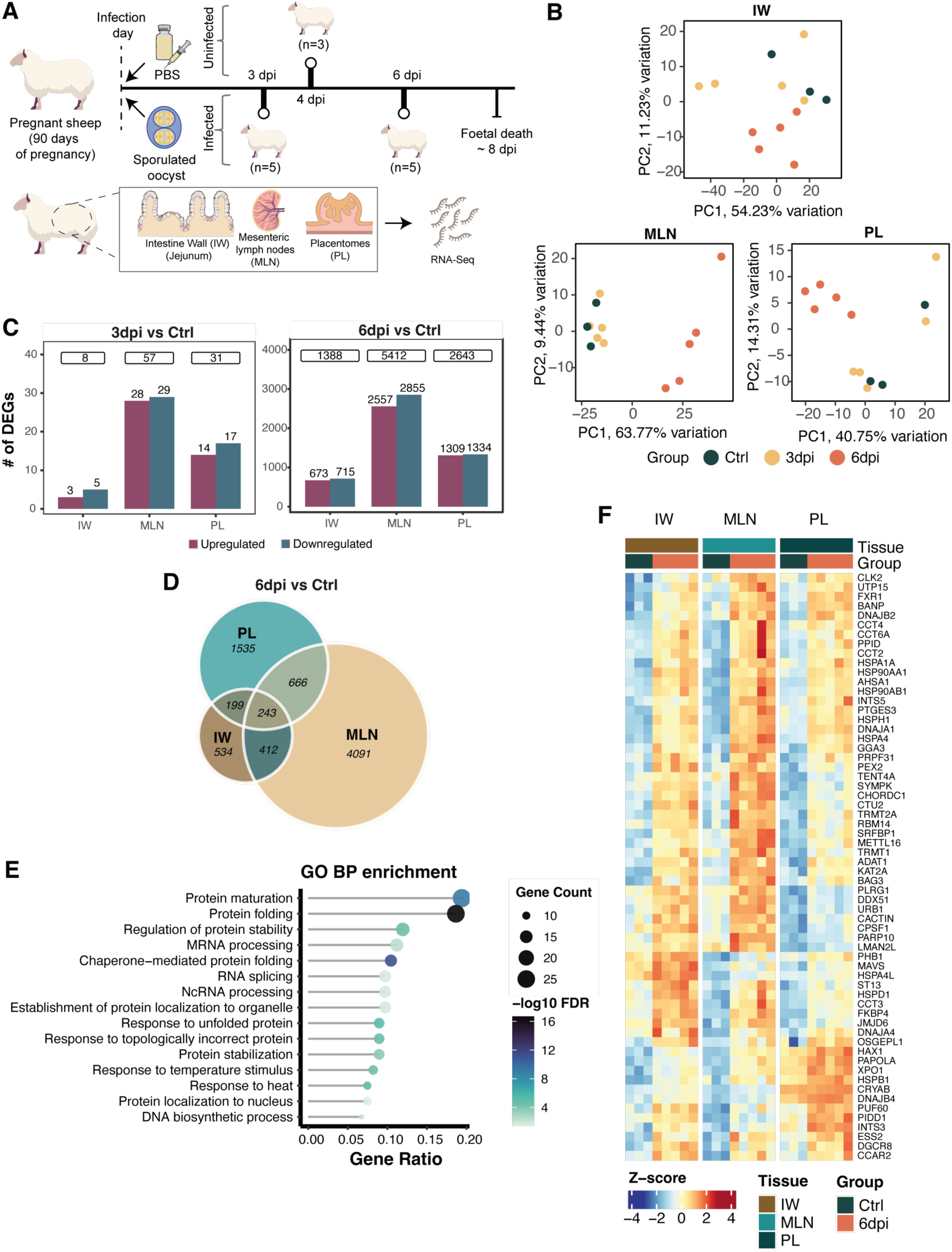
**Transcriptomic signatures of *T. gondii* infection in pregnant sheep**. A. Diagram of the experimental design for *T. gondii* infection in pregnant sheep. B. PCA of the transcriptome profile of *T. gondii-*infected and control samples, separated by tissue, using the top 1,000 most variable genes in each tissue. Colors indicate experimental condition (green for control samples, yellow for 3 dpi infected samples, and orange for 6 dpi infected samples). Percentages of variance explained by PC1 and PC2 are shown on the respective axes. C. Bar plots show the number of upregulated (red) and downregulated (blue) DEGs in each tissue, comparing infected samples at 3 dpi (left) and 6 dpi (right) with control samples. The number of up- and downregulated DEGs is indicated above each corresponding bar, and the total number of DEGs is shown in a box above both bars. D. Venn diagram showing overlapping DEGs between infected samples from 6 dpi and control samples in IW, MLN and PL. E. Dot plot showing GO enrichment analysis of commonly upregulated DEGs at 6 dpi. Dot color represents the -log10(FDR), and dot size indicates the number of DEGs associated with each term. F. Heatmap showing the expression (Z-score) of commonly upregulated DEGs at 6 dpi associated with GO enriched terms from E. Column colors indicate tissue and experimental condition, with green representing controls and orange representing 6 dpi infected samples.

Principal Component Analysis (PCA) of all samples revealed a clear separation primarily according to tissue identity (Fig. S1A). In contrast, tissue-specific PCA showed distinct clustering of infected samples at 6 dpi, whereas control and 3 dpi samples did not segregate distinctly by infection status (Fig. 1B). To quantify the degree of overlap in transcriptional variability across tissues and stages of infection, we next examined the top 1,000 most variable genes used for PCA in each tissue. Only 86 genes were shared among all three tissues, highlighting the marked tissue specificity of the host transcriptomic response (Fig. S1B). In agreement with the PCA results, hierarchical clustering of samples based on their gene expression recapitulated the dominant tissue-driven structure of the data and further demonstrated a clear separation of infected samples at 6 dpi within each tissue compartment (Fig. S1C).

To investigate the host transcriptomic changes induced by *T. gondii* infection, we performed a differential expression analysis comparing data from IW, MLN, and PL between infected (3 dpi or 6 dpi) and control sheep (Table S1). At 3 dpi, transcriptional changes between infected and control samples were limited, with 8, 57, and 31 differential expressed genes (DEGs; False Discovery Rate, FDR < 0.05) detected in IW, MLN, and PL tissues, respectively, with no DEGs shared between tissues (Fig. 1C, Fig. S2A). The MLN, which exhibited the strongest transcriptional response at this stage, showed differential expression of some immune-related genes, including downregulation of *TNFSF11*, *CTSC*, *ALOX5*, *APOA1*, *KLRD1*, and *CXCR4*, and upregulation of *NCR3* in infected animals (Table S1).

At 6 dpi, the transcriptional response expanded markedly, with 1,388, 5,412, and 2,643 DEGs identified in IW, MLN, and PL, respectively (Fig. 1C). Of these, 243 DEGs were shared across all tissues, of which 178 were consistently upregulated and 56 were consistently downregulated (Fig. 1D, Fig. S2B). Over-representation analysis of the commonly upregulated DEGs revealed a significant enrichment of GO biological processes (FDR < 0.05), with the 15 terms showing the highest gene ratio predominantly related to protein maturation and folding, chaperone activity, and response to temperature (Fig. 1E, Table S2). Among the genes associated with these biological processes are several encoding chaperones, including *HSPA1A*, *HSPB1*, *HSPH1*, *HSP90AA1*, *HSP90AB1*, and *HSPA4*, as well as co-chaperones such as *DNAJA1*, *DNAJB2*, and *CHORDC1*. Additional genes such as *FKBP4*, and *BAG3*, which interact with chaperones and contribute to proteostasis, were also upregulated in infected samples (Fig. 1F). In contrast, no biological processes were significantly enriched among the commonly downregulated DEGs.

### *Toxoplasma gondii* infection promotes a tissue-specific immune response at 6 days post-infection

To characterize tissue-specific host responses at 6 dpi, we performed Gene Set Enrichment Analysis (GSEA) using Hallmark gene sets (Fig. 2A, Table S3).

**Figure 2.**
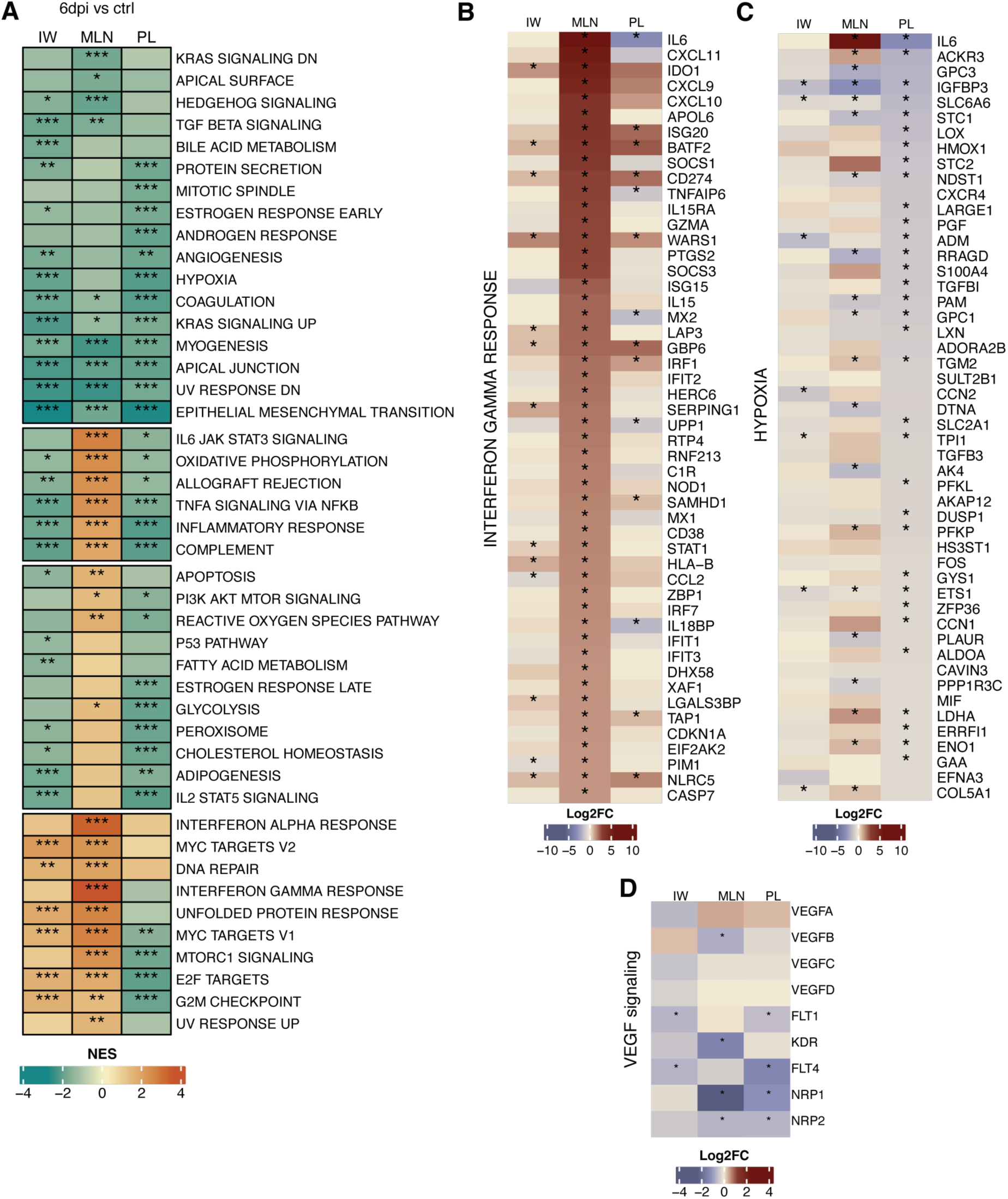
**GSEA highlights pathways modulated during *T. gondii* infection in pregnant sheep**. A. Heatmap of GSEA results for each tissue (columns) using the Hallmark gene set, shown separately for 3 dpi and 6 dpi. The color gradient represents normalized enrichment scores (NES), ranging from negative (blue, pathway inhibition in infected samples) to positive (orange, pathway activation in infected samples). Asterisks indicate statistically significant enrichments (*** FDR < 0.001, **FDR < 0.01, * FDR < 0.05). B-D. Heatmap shows the top 50 genes with the highest log2FC in MLN associated with interferon gamma response (B), top 50 genes with lowest log2FC in PL associated to hypoxia gene set (C), and the differential expression of VEGF signaling-related genes, including VEGF receptors and co-receptors (D), comparing infected samples at 6 dpi with controls. Log2FC values for 6 dpi versus control are shown across the three tissues. The color gradient represents log2FC, ranging from blue (downregulated) to red (upregulated). Asterisks denote DEGs (FDR < 0.05).

In the IW, upregulated pathways were mainly associated with cell cycle progression, proliferation, and cellular stress responses, including MYC targets, E2F targets, G2/M checkpoint, DNA repair, and unfolded protein response pathways. Several of these pathways were also enriched in the MLN, whereas cell cycle-related pathways were downregulated in the PL, indicating suppression of proliferative and cell cycle-associated programs in this tissue.

The MLN showed the strongest immune response with upregulation of inflammatory response, TNFα signaling via NF-κB, IL6-JAK-STAT3 signaling, complement and allograft rejection, and oxidative phosphorylation, many of which were downregulated in both the IW and the PL. Notably, IFN-γ and IFN-α response pathways were exclusively upregulated in the MLN, revealing a robust interferon-driven immune response restricted to lymphoid tissue. This signature was primarily driven by increased expression of *IL6*, *CXCL11*, *IDO1*, *CXCL9*, *CXCL10*, and *APOL6* in infected samples (Fig. 2B). Although interferon response pathways were not significantly enriched in the IW and PL, several interferon-stimulated genes (ISGs), including *IDO1*, *ISG20*, *BATF2*, *CD274*, *WARS1*, *GBP6*, and *NLRC5*, were upregulated in these tissues.

Strikingly, the PL exhibited a globally repressed transcriptional profile, with all significantly enriched Hallmark gene sets showing negative normalized enrichment scores (NES < 0). Alongside cell cycle and immune-related pathways, hormone responses, angiogenesis, hypoxia, glycolysis and peroxisome pathways were downregulated (Fig. 2A). Reduced expression of *PGF*, *ACKR3* and *CXCR4*, together with the endothelial regulator *ETS1* and hypoxia related genes *SLC2A1* and *HMOX1*, suggested altered vascular and hypoxia-related signaling (Fig. 2C). Genes involved in extracellular matrix remodeling and cell-matrix interactions, including *LOX*, *TGFB1*, *LARGE1* and *S100A4*, were also downregulated, pointing to changes in placental tissue remodeling. Given the reduced expression of *PGF*, a member of vascular endothelial growth factor (VEGF) family, which plays a central role in angiogenesis (21), we further examined the expression of key VEGF signaling components (Fig. 2D). This analysis revealed downregulation of VEGF receptors (*FLT1* and *FLT4*) and co-receptors (*NRP1* and *NRP2*) in placentomes from infected samples, suggesting impaired VEGF-mediated angiogenic signaling in infected placental tissues. Notably, some of these genes were also downregulated in the MLN and IW, indicating that modulation of VEGF signaling may occur across multiple tissues during acute infection.

### *Toxoplasma gondii* infection differentially modulates immune and angiogenesis regulatory programs across tissues

To extend our characterization beyond enrichment analysis, we estimated pathway activity scores using the PROGENy database (22) implemented in decoupleR (23), employing the statistic value from differential expression analysis as input. At 6 dpi, the MLN exhibited strong activation of JAK-STAT, MAPK, NF-κB, and PI3K pathways, whereas JAK-STAT and MAPK pathways were moderately activated in the IW and PL, with PI3K pathway suppressed in the PL and NF-κB pathway inhibited in the IW. In the PL, androgen, and TNFα pathways were repressed, while p53 and VEGF pathways were activated. Across all tissues, hypoxia, TRAIL, and TGF-β pathways were inhibited (Fig. 3A).

**Figure 3.**
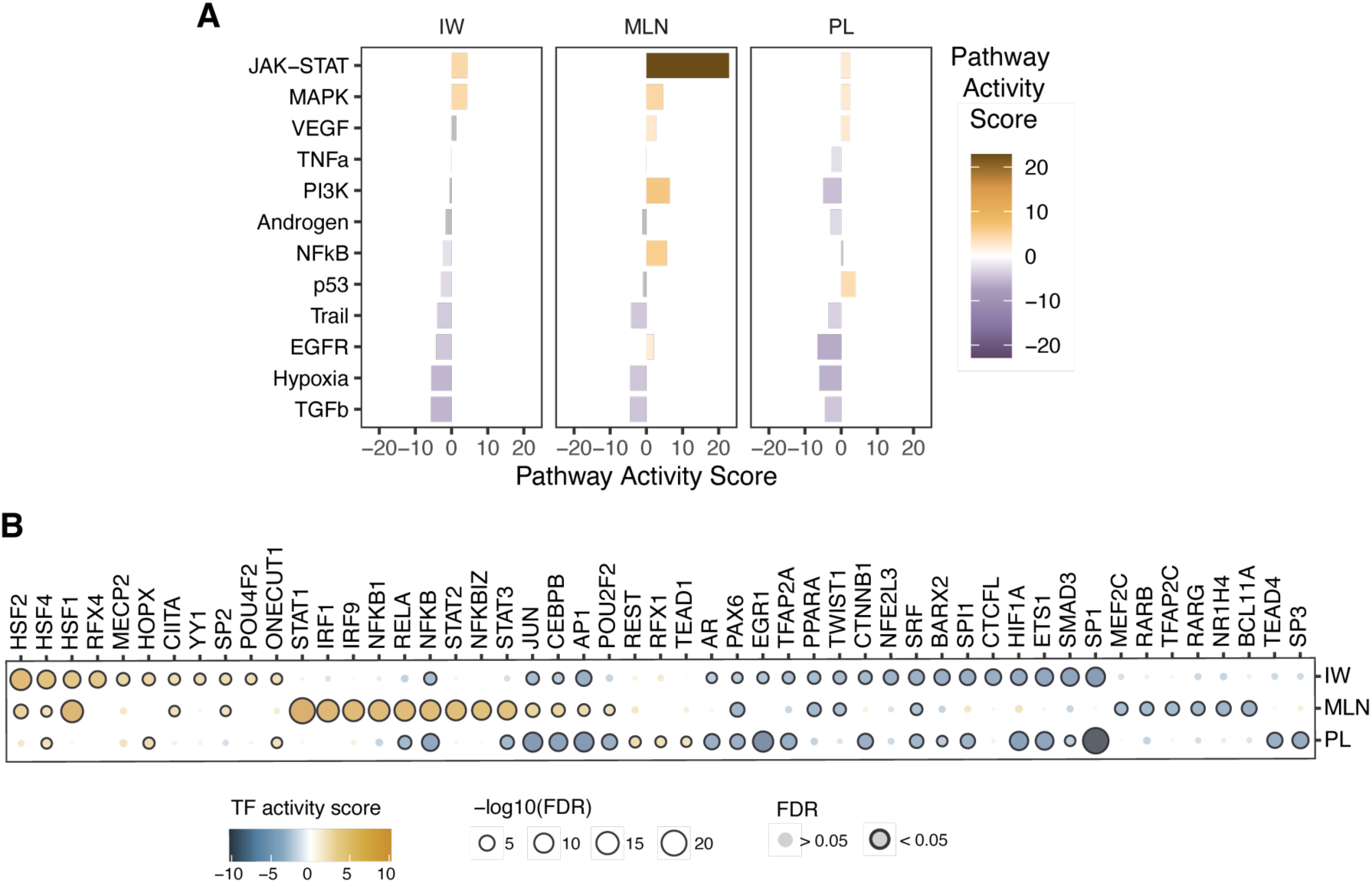
Pathways and Transcription factor activity across tissues following *T. gondii* infection at 6 dpi. **A**. Pathway activity scores inferred with PROGENy via decoupleR. Bar color and x-axis indicate activity, from purple (higher in controls) to orange (higher in infected samples). Gray bars indicate non-significant pathways (P > 0.05). **B**. TF activity scores inferred using CollecTRI via decoupleR. The top 30 significant TFs (FDR < 0.05) per tissue are shown. Dot size indicates -log10(FDR) with TFs showing significant differential activity outlined in black. Color gradient represents TF activity score, from blue (higher in controls) to yellow (higher in infected samples).

Next, we investigated the transcriptional regulators underlying these tissue-specific responses at 6 dpi by inferring Transcription Factor (TF) activity using decoupleR together with the CollecTRI gene regulatory network framework (24). Across all tissues, infection was consistently associated with increased activity of heat-shock factor (HSF) family members, in agreement with the induction of heat-shock protein genes upon infection (Fig. 3B).

Beyond this shared stress response, infection elicited highly tissue-specific regulatory programs. In the MLN, infection induced a strong activation of immune and interferon-associated transcriptional networks, characterized by increased activity of STAT family members (STAT1, STAT2, STAT3), interferon regulatory factors (IRF1, IRF2, IRF7, IRF9), and key components of the NF-κB pathway (NFKB1, NFKB2, RELA, REL). This regulatory signature is consistent with a robust inflammatory and interferon-driven response upon infection in lymphoid tissue. In contrast, infection in both the IW and PL was not accompanied by activation of these canonical immune TFs but instead showed a coordinated reduction in the activity of endothelial cell TFs, including HIF1A, SMAD3, ETS1, CTNNB1, and SP1 (25). Notably, the PL displayed a distinct induced regulatory program characterized by increased activity of REST, RFX1, and TEAD1, together with reduced activity of SP3 and TEAD4.

### Temporal accumulation of NK cells and M1 macrophages in the mesenteric lymph node during *Toxoplasma gondii* infection

To characterize immune cell infiltration across tissues, CIBERSORTx (26) was applied to bulk RNA-seq data using the LM22 signature matrix as reference. The resulting absolute immune score varied across tissues and experimental conditions (Fig. 4A). Notably, the PL exhibited the lowest levels of immune cell infiltration among all tissues analysed. In contrast, moderate immune infiltration was observed in the IW and, as expected, the highest immune cell abundance was observed in the MLN. Among all tissues, the MLN was the only one showing significant differences in immune cell abundance among experimental conditions, with significantly higher levels at 6 dpi compared with both the control (*p* = 0.007) and 3 dpi samples (*p* = 0.019) (Fig 4A). In MLN, patterns of immune cell proportions were consistent across samples within each experimental condition (Fig. 4B). Comparison of cell subsets revealed higher proportions of activated NK cells and M1 macrophages at 6 dpi than in both controls and 3 dpi samples (*p* = 0.0191 and *p* = 0.0094, respectively, for both cell types). Conversely, memory B cells decreased between 3 and 6 dpi (*p* = 0.0074), as did M0 macrophages (*p* = 0.0147) (Fig. 4C).

**Figure 4.**
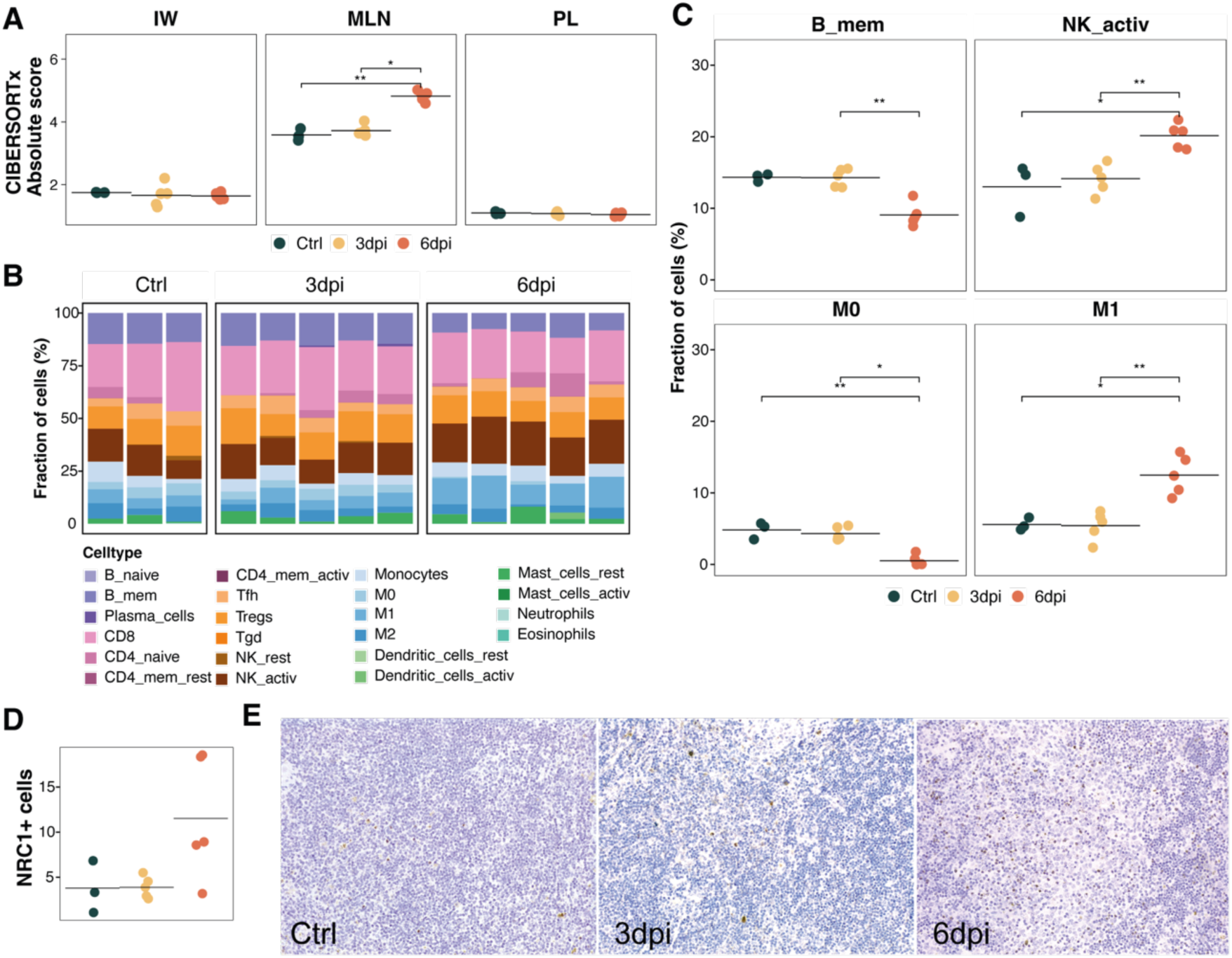
Immune cell composition inferred by deconvolution analysis. A. Dot plots showing absolute abundance scores of inferred immune cell types, separated by tissue. Dot color indicate experimental condition (green for control samples, yellow for 3 dpi infected samples, and orange for 6 dpi infected samples). B. Bar plots representing the relative proportion (%) of each immune cell type per sample. Colors indicate different cell types. C. Dot plots showing the percentage of immune cell types in MLN with significant proportional changes between experimental conditions. Asterisks indicate statistically significant pairwise differences based on Dunn’s post hoc test following the Kruskal-Wallis test (** P < 0.01, * P < 0.05). D. Dot plot depicting the mean number of NK cells (NCR1⁺) in placentomes across experimental conditions. E. Immunohistochemical detection of NCR1-positive NK cells in the MLN. Positive staining is evident as brown cytoplasmic immunolabelling in scattered small mononuclear cells distributed throughout the paracortical and medullary areas of the lymph node. IHC, 200×.

To complement the transcriptomic analysis, immunohistochemistry (IHC) was used to characterize immune cell populations in MLNs. NRAMP1 staining was concentrated in follicles, whereas NK cells were predominantly localized in the medulla. TNF-α staining was more widespread, with higher abundance in the medulla and germinal centres. Although NRAMP1 and TNF-α immunolabelling did not differ between groups, NCR1⁺ cell abundance tended to increase at 6 dpi, consistent with the transcriptomic data, but this difference did not reach statistical significance (*p* = 0.09) (Fig 4D, E).

### *Toxoplasma gondii* infection downregulates cell cycle and extracellular matrix organization programs while promoting stress-associated transcriptional responses in placentomes

To characterize placental transcriptional dynamics during infection, 18,364 expressed genes were grouped into six clusters based on their expression profiles in control and infected animals (Fig. 5A, B, Table S4). Over-representation analysis of Hallmark gene sets and Reactome pathways was performed to identify biological processes potentially involved in the molecular mechanisms leading to early fetal death following *T. gondii* infection (Fig. 5C, Fig. S3, Table S4).

**Figure 5.**
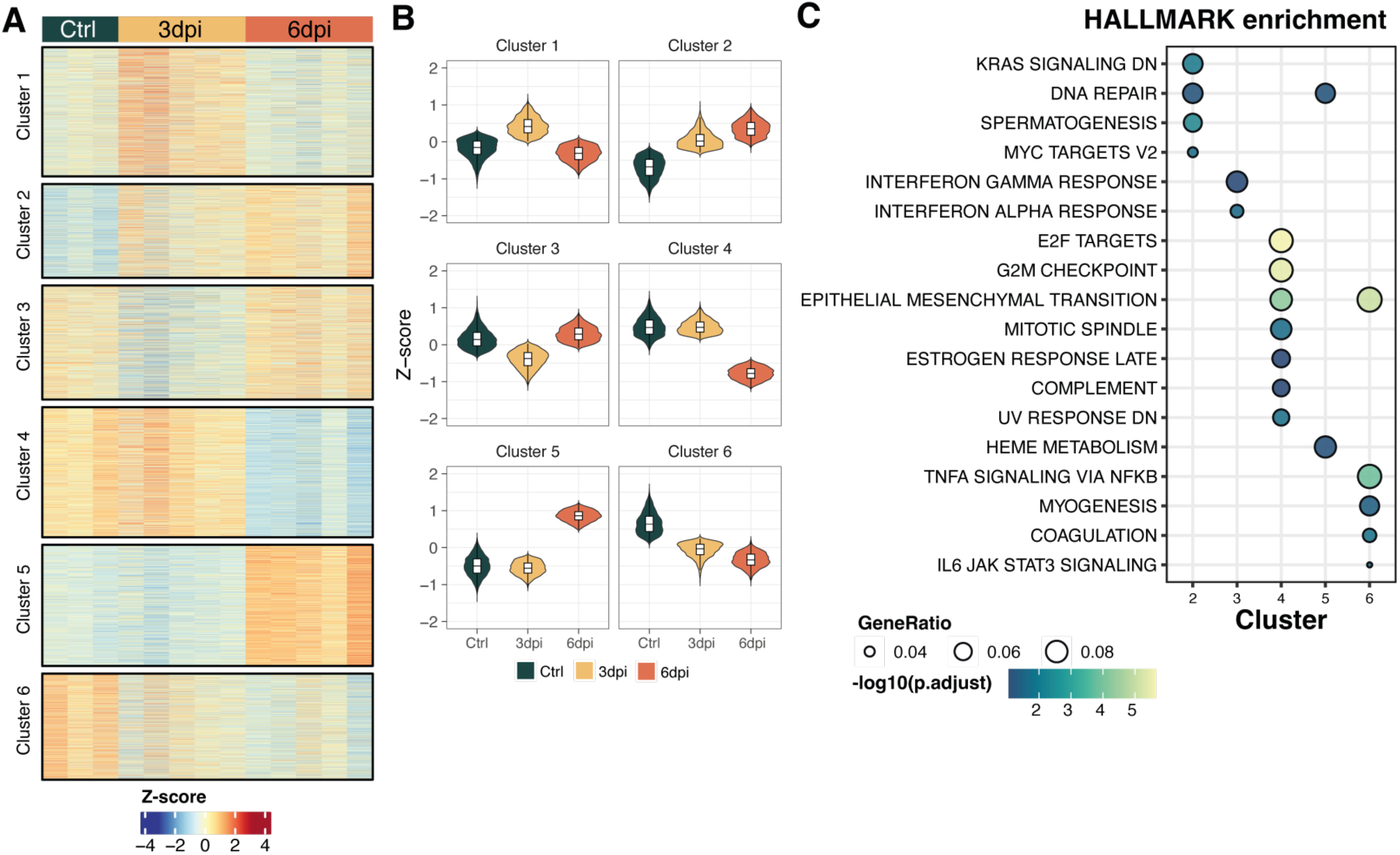
Gene expression dynamics during early *T. gondii* infection in placentomes. A. Heatmap showing the Z-score-normalized expression of genes in each cluster across control and *T. gondii*-infected samples. The color gradient represents standardized gene expression levels (Z-scores). The top annotation indicates the experimental condition (green for control samples, yellow for 3 dpi infected samples, and orange for 6 dpi infected sample). B Violin plots showing the distribution of mean gene expression (Z-scores) for each cluster under each experimental condition. C. Over-Representation analysis of Hallmark gene sets. The 15 most significant terms for each cluster (FDR < 0.1) are shown. Dot size represents the gene ratio, and dot color represents -log10(FDR).

Two clusters (2 and 5) showed increasing expression during infection, although they differed in the timing and functional features of their responses (Fig. 5, Fig. S3). Both were enriched in DNA repair-related pathways, while cluster 2 showed a progressive increase associated with genome maintenance, chromatin modification, RNA processing and protein ubiquitination. Cluster 5, in contrast, was predominantly upregulated at 6 dpi and was enriched in cellular stress-response pathways, including HSF1 activation and TP53 regulation. The opposite pattern was observed in clusters 4 and 6, which showed progressively decreasing expression (Fig. 5A, B). Cluster 4 was associated with cell cycle regulation, mitosis, epithelial-mesenchymal transition (EMT) and complement, whereas cluster 6 was enriched in extracellular matrix organization and remodeling, coagulation and inflammatory signaling pathways, including TNFα signaling via NF-κB and IL6- JAK-STAT3 signaling (Fig. 5C, Fig. S3). Finally, cluster 3, enriched in interferon response, showed a transient decrease in expression at 3 dpi, followed by an increase at 6 dpi to levels slightly above those observed in controls (Fig. 5).

## DISCUSSION

This study characterizes tissue-specific transcriptional responses to *T. gondii* infection in pregnant sheep during the acute phase preceding early fetal death (4). Our findings reveal dynamic, tissue-specific responses that provide insight into molecular events potentially associated with early reproductive failure.

At 3 dpi, transcriptional responses were limited across tissues, consistent with the low or undetectable parasite burden (8). Nevertheless, the MLNs showed early immune-related transcriptional changes, predominantly involving gene downregulation, consistent with previously described immune evasion strategies of *T. gondii* (27, 28). At 6 dpi, transcriptional changes were more pronounced, particularly in the MLNs, where parasite burden was highest (8). Although most DEGs were tissue-specific, 243 were shared across all three tissues and enriched in chaperone activity, proteostasis and temperature response, pointing to a conserved cellular stress response during early infection. This signature included upregulation of *HSPA1A*, *HSPA4* and *HSPB1*, coinciding with the peak of fever (8), and consistent with previous reports of HSP induction during *T. gondii* in vivo and in vitro infections (28–30). Elevated expression of HSP70 inhibits apoptosis in *T. gondii*-infected cells, promoting host cell survival while favoring parasite intracellular replication and persistence (31, 32). Thus, chaperone upregulation may reflect a dual response that mitigates host cellular stress while creating conditions favourable to the parasite, which can exploit endoplasmic reticulum stress pathways to maintain proteostasis, modulate host cell survival and promote its dissemination (33–35).

At 6 dpi, the IW showed a predominant suppression of proinflammatory immune pathways, such as TNF-α signaling via NF-κB, IL-2-STAT5 signaling, and pathways linked to the complement system and the inflammatory response. This contrasts with murine models, in which *T. gondii* induces severe ileitis from 8 dpi, driven by an exaggerated Th1 response and excessive production of proinflammatory mediators (20, 36, 37). Although digestive tract lesions have been reported in several infected animal species following *T. gondii* infection (38), such inflammation was not observed in sheep. However, given the extensive length of the ovine small intestine, examination of additional intestinal sections may be needed to fully assess pathology.

In the MLN, infection induced a shift from an early suppressed inflammatory environment (3 dpi) to a strong proinflammatory response at 6 dpi, marked by upregulation of *IFNG*, *TNF*, which are critical for controlling *T. gondii* infection (39, 40), alongside *CXCL11*, *IL6*, *CXCL9* and *CXCL10*, indicating a robust activation of interferon- and cytokine-mediated immune pathways in this tissue, as also observed in pigs (41). Our data also revealed activation of JAK-STAT and NF-κB signaling pathways, which mediate interferon signaling and inflammatory responses against intracellular pathogens (42–44) and regulate inflammatory responses during type II *T. gondii* infection (45, 46). Increased activity of STAT, IRF and NF-κB transcription factors further supports the involvement of these pathways in coordinating the MLN immune response during acute infection.

Consistent with immune activation, deconvolution analysis revealed expansion of immune cell populations in the MLN at 6 dpi, consistent with the observed adenomegaly (5, 8). Infected samples showed increased proportions of activated NK cells and M1 macrophages, with NK cell expansion further confirmed by IHC. Similar findings have been reported in other *T. gondii* infection models (47, 48). These innate immune cells are major sources of IFN-γ and contribute to protective Th1 responses against intracellular pathogens (49–52). Accordingly, NK cell depletion or dysfunction impairs Th1 responses and increases susceptibility to *T. gondii* (53). Our results provide direct evidence in sheep that NK cells constitute a key component of the early immune defense against *T. gondii*.

The placenta emerges as a critical target tissue during early fetal death, as placental thrombosis is the only lesion described in the absence of parasite invasion (7, 8). Despite undetectable parasite DNA, placentomes showed upregulation of ISGs, as *ISG20*, *IRF1*, *BATF2* and *GBP6*, at 6 dpi, suggesting an early immune activation. Despite this immune stimulation, inflammatory response was suppressed, alongside downregulation of angiogenesis and hypoxia signaling, both essential for placental homeostasis (54, 55). Reduced expression of the VEGF receptors (*FLT1* and *FLT4*), co-receptors (*NRP1* and *NRP2*), and *PGF*, together with downregulation of PI3K signaling, suggests impaired VEGF-mediated vascular signaling and placental remodeling, potentially contributing to fetal loss (56–59). Upregulated IFN-induced chemokines (*CXCL9*, *CXCL10* and *CXCL11)* in the MLN may also contribute to angiogenic suppression through their antiangiogenic activity (60–62). Also, hypoxia signaling and HIF-1α activity were reduced at 6 dpi, contrasting with their activation in preeclampsia (63, 64). Although HIF-1α promotes angiogenesis, it can also induce antiangiogenic factors (sFlt-1 and sEng) and suppress PGF, impairing endothelial function and trophoblast invasion (65–67). Finally, our data also revealed downregulation of genes involved in cell cycle regulation and extracellular matrix organization, consistent with previous evidence that *T. gondii* disrupts E2F activity and mitotic regulators, promoting G2/M arrest (27). In placenta, this suppression of E2F targets and G2/M checkpoint machinery indicates a severe proliferative blockade in trophoblast populations (68, 69). These alterations may impair trophoblast proliferation and placental remodeling, compromise the maternal-fetal interface and potentially contribute to pregnancy loss (69, 70), although further studies are needed to establish their functional consequences.

In summary, acute ovine toxoplasmosis elicited contrasting tissue-specific responses, with restrained intestinal inflammation, strong proinflammatory activation in MLNs and placental changes suggestive of impaired angiogenesis and tissue remodeling despite undetectable parasite invasion. These alterations may compromise placental function and contribute to fetal loss, although the cellular heterogeneity of the placenta limits the interpretation of bulk transcriptomic data. Resolving cell-specific responses will be essential to understand how systemic immune activation and local placental dysregulation contribute to reproductive failure during acute ovine toxoplasmosis. Ultimately, unraveling the mechanisms of early fetal death in the ovine model provides an essential translational framework to explore human congenital toxoplasmosis and identify diagnostic biomarkers and therapeutic targets for early pregnancy losses.

## MATERIALS AND METHODS

### Experimental design and sampling

The experimental design was described by Sánchez-Sánchez et al. (8). Briefly, thirteen pregnant sheep (18 months old) were included. At 90 days of gestation, ten animals were inoculated with 10^3^ sporulated oocysts of *T. gondii* isolate TgShSp1, while three control sheep were inoculated with 50 ml of phosphate-buffered saline (PBS) at the same time. Infected sheep were euthanized at 3 (n = 5) and 6 (n = 5) dpi, and control animals at 4 dpi. Medial jejunal mesenteric lymph nodes (MLNs), jejunal intestinal wall (IW), and placental samples (pools of four placentomes from the intermediate uterine region) were collected for RNA sequencing (RNA-seq) and stored at -80 °C in RNAlater (Thermo Fisher Scientific, Paisley, UK) until used. Total RNA was extracted using the Maxwell® 16 LEV simplyRNA Purification Kit, developed for the automated Maxwell® 16 System (Promega, Madison, WI, USA), following the manufacturer’s recommendations. Representative MLN specimens were collected and immediately fixed in 10% neutral buffered formalin. After fixation, samples were trimmed, dehydrated through a graded series of alcohols, cleared in xylene and embedded in paraffin wax using standard histological procedures.

### RNA-seq library preparation and sequencing

Total RNA concentration and integrity were assessed using the Qubit® RNA BR Assay Kit (Thermo Fisher Scientific) and the RNA 6000 Nano Assay on an Agilent 2100 Bioanalyzer. RNA-seq libraries were generated using the KAPA Stranded mRNA-Seq Kit for Illumina® platforms (Roche Diagnostics, Indianapolis, IN, USA), following the manufacturer’s protocol. Briefly, 500 ng of total RNA was used as input. mRNA was selectively enriched through oligo-dT magnetic bead capture of the poly-A tail, followed by enzymatic fragmentation. Strand specificity was maintained through second-strand synthesis using dUTP. The resulting double-stranded cDNA was end-repaired, 3′ adenylated, and ligated to Illumina-compatible adapters containing unique dual indices and unique molecular identifiers (Integrated DNA Technologies, Coralville, IA, USA). The adapter-ligated libraries were amplified with 15 cycles of PCR. Final library quality and fragment size distribution were verified using the DNA 7500 assay on the Agilent 2100 Bioanalyzer. Libraries were sequenced on a NovaSeq 6000 (Illumina, San Diego, CA, USA) with 2×100 bp reads and ∼30 million paired-end reads per sample, using dual indexing. Data processing, including base calling and quality scoring, was done with RTA v3.4.4, and FASTQ files were generated.

### RNA-seq preprocessing

Quality control of RNA-seq reads was performed using FastQC (v0.11.4) (71), and high-quality reads were then aligned to the *Ovis aries* reference genome (ARS-UI_Ramb_v2.0; ENSEMBL release 111), using STAR (v2.7.9a) (72). Gene-level read counts were generated using the GeneCounts option implemented in STAR.

### Differential expression and functional enrichment analysis

Differential expression analysis was performed using the DESeq2 R package (v1.42.1) (73). Genes with mean raw counts below 1 were excluded during exploratory analysis, and variance-stabilizing transformation (VST) was applied for principal component analysis (PCA) of the 1,000 most variable genes using PCAtools (v2.14.0) (74). Within each tissue, infected samples at 3 and 6 dpi were compared separately with uninfected controls. Genes with fewer than 10 counts in at least three samples were filtered out, and differential expression was assessed using the DESeq2 Wald test, with False Discovery Rate (FDR) < 0.05 defining differentially expressed genes (DEGs). For downstream analysis, ENSEMBL gene IDs from sheep were mapped to human orthologues using the biomaRt package (v2.61.3) (75).

Functional enrichment was performed using *clusterProfiler* R package (v4.10.1) (76). Over- representation analysis of shared DEGs across tissues used Gene Ontology (GO) biological process (77, 78), with up- and downregulated genes analyzed separately against all expressed genes as the background. Terms containing at least three genes and meeting FDR < 0.05 were considered significant. Gene Set Enrichment Analysis (GSEA) (79) was performed using the Hallmark gene sets from Molecular Signature Database (MSigDB) (80), accessed via the msigdbr R package (v7.5.1) (81) for human data. Genes were ranked by DESeq2 Wald statistics, and gene sets with FDR < 0.05 were considered significantly enriched.

### Transcription Factor regulon and pathway activity analysis

Transcription Factor (TF) activity was inferred using the decoupleR R package (v2.8.0) (23), using the CollecTRI gene regulatory network (24). TF activity was estimated using the univariate linear model based on DESeq2 Wald statistics, considering TFs with at least five target genes. Significant TFs were identified using a Benjamini-Hochberg-adjusted FDR threshold of 0.05. Pathway activity was inferred using PROGENy (22), and the multivariate linear model (run_mlm), considering the top 500 genes per pathway (minimum five), with *P* values < 0.05 were considered significant.

### Gene expression clustering and pathway enrichment analysis in placentomes

Expressed genes were clustered according to their expression profiles across control and infected samples. VST expression values were transformed to Z-scores, and partitioning around medoids (PAM) was applied to mean expression values for each experimental condition. The optimal number of clusters was determined by evaluating average silhouette widths and biological interpretability. Functional enrichment was conducted using the *compareCluster()* function from the clusterProfiler (v4.10.1) (76) against Hallmark and Reactome gene sets from MSigDB (80), retrieved using msigdbr (v7.5.1) (81). Terms with at least three genes and FDR < 0.1 were considered significant.

### Cell type deconvolution of bulk RNA-Seq

Immune cell composition in bulk RNA-seq samples was estimated using CIBERSORTx (26). Expression values were normalized to transcripts per million (TPM), and sheep genes were converted to human orthologues. Deconvolution was performed using the LM22 leukocyte signature matrix (82), with B-mode batch correction, 100 permutations, and absolute mode. Differences between conditions were assessed using the non-parametric Kruskal-Wallis test, followed by Dunn’s post hoc test for pairwise comparisons.

### Immunohistochemistry (IHC)

Immunohistochemical labelling was performed on formalin-fixed, paraffin-embedded (FFPE) sections (3 μm) of medial jejunal MLNs. Sections were first deparaffinised in xylene and rehydrated through a graded series of ethanol. For NRAMP1 detection, enzymatic epitope retrieval was carried out using Trypsin Antigen Retrieval Solution (ab970, Abcam, Cambridge, UK). For NK cell and TNF-α detection, heat-induced epitope retrieval (HIER) was performed for 20 min at 96 °C using Target Retrieval Solution (Agilent/Dako, Santa Clara, CA, USA) at pH 9 (Tris/EDTA buffer, S2367) and pH 6 (citrate buffer, S2369), respectively.

Endogenous peroxidase activity was blocked by incubating the sections in methanol containing 3% hydrogen peroxide for 30 min in the dark. Sections were subsequently incubated overnight at 4°C with the following primary antibodies diluted in Dako Antibody Diluent (K8006, Agilent/Dako): rabbit anti-NRAMP1 (1:50, sc-20113, Santa Cruz Biotechnology Inc., Dallas, TX, USA), mouse anti-NK cells (1:50, MCA5933GA, Bio-Rad Laboratories, Hercules, CA, USA), and mouse anti-TNF-α (1:100, MCA2334, Bio-Rad Laboratories).

After rinsing, the EnVision System-HRP (K4003 for NRAMP1 and K4001 for NK and TNF-α; Agilent/Dako) was applied for 40 min at room temperature in the dark. Immunoreactivity was visualised using 3,3′-diaminobenzidine (DAB) chromogen (K3468, Agilent/Dako). Finally, sections were counterstained with haematoxylin, dehydrated through graded alcohols, cleared in xylene, and mounted with DPX medium.

### Image analysis

Quantification of DAB-positive immunolabeling was performed on 20 randomly selected fields per sample, captured at 20× magnification, using a custom macro developed in ImageJ (83). Briefly, colour deconvolution ("H DAB" vector) separated haematoxylin and DAB signals; manually thresholded tissue and DAB+ masks were combined by logical AND, and the percentage of immunomarked area was calculated as the ratio of DAB-positive to total tissue area per field.

### Statistical analysis and data visualization

All statistical analyses were performed in R (v4.3.2) (84). Statistical differences in the numbers of NCR1, TNFα and NRAMP1-positive cells assessed by IHC, as well as in immune cell fractions estimated by deconvolution, across experimental conditions were evaluated using the non-parametric Kruskal-Wallis test, followed by Dunn’s post hoc test for pairwise comparisons between groups. For IHC analyses, the mean value across the 20 fields examined was calculated for each animal.

Heatmaps and upset plots were generated using the ComplexHeatmap (v2.25.2) (85) and ComplexUpset (v1.3.6) (86, 87) R packages, respectively. Additional data visualizations were created with ggplot2 (v3.5.1) (88).

## Data availability

The raw sequencing data generated in this study has been deposited in the European Nucleotide Archive (ENA) under the accession number PRJEB105728 (https://www.ebi.ac.uk/ena/browser/view/PRJEB105728).

## Acknowledgments

We thank the SCBI (Supercomputing and Bioinformatics) Center of the University of Málaga for providing the computational resources used for part of the transcriptomic data preprocessing. This work was supported by grant PID2022-138673OB-C21, funded by the MICIU/AEI/ 10.13039/501100011033 and by ERDF/EU; and by the Community of Madrid, Spain (TEC- 2024/BIO-66/SALAINDEC-CM).

